# Optimized MALDI-TOF mass spectrometry enables reliable identification of freshwater snails from schistosomiasis-endemic areas in Mauritania

**DOI:** 10.64898/2026.06.01.729200

**Authors:** Lemat Nakatt, Lionel Almeras, Ouldabdallahi Mohamed Moukah, Adama Zan Diarra, Ali Ould Mohamed Salem Boukhary, Stéphane Ranque

## Abstract

Freshwater snails act as intermediate hosts for parasites affecting both humans and livestock, including schistosomes. In Mauritania, however, the diversity, distribution, and infection status of these snails remain poorly documented. This study aimed to identify freshwater snail species collected from two schistosomiasis-endemic areas in southern Mauritania, characterize their spatial distribution, and assess their infection rates using molecular tools Malacological surveys were conducted in Kankossa and Oued Rawdha during the 2023 rainy season. A total of 806 snail specimens were collected and preserved in 70% ethanol at 4 °C prior to analysis. Five species were identified morphologically and confirmed by molecular analysis: *Bulinus truncatus*, *B*. *forskalii*, *B*. *senegalensis*, *B*. *umbilicatus*, and *Melanoides tuberculata*. MALDI-TOF MS generated high-quality spectra for 99.0% of specimens and correctly identified 99.9% of analyzable samples after molecular confirmation of discrepant cases. Preservation in ethanol at 4 °C markedly improved spectral quality compared with previously reported room-temperature storage conditions

Distinct ecological distributions were observed according to water body type. *B. senegalensis* and *B. umbilicatus* were exclusively collected from temporary ponds, whereas *B. truncatus*, *B. forskalii*, and *M. tuberculata* were found in permanent water bodies. Real-time PCR screening detected *Schistosoma haematobium* complex DNA in 239/798 (29.9%) specimens, with substantially higher infestation rates in Kankossa than in Oued Rawdha.

These findings demonstrate that MALDI-TOF MS is a rapid, accurate, and field-compatible tool for freshwater snail identification, including closely related species that are difficult to distinguish morphologically. This approach could facilitate large-scale epidemiological surveillance and improve monitoring of schistosomiasis transmission dynamics in endemic settings.

## Introduction

Accurate identification of freshwater snail species and associated pathogens is essential for designing effective schistosomiasis control program (Gryseels *et al*., 2006; Abe *et al*., 2018). For decades, the identification of medically and veterinary important freshwater snails has relied on macroscopic and microscopic morphological criteria and/or molecular methods (Raahauge & Kristensen, 2000; Kane et al., 2008; Zein-Eddine et al., 2014; Hamlili et al., 2021).

Matrix-assisted laser desorption/ionization time-of-flight mass spectrometry (MALDI-TOF MS), a novel proteomic method, has emerged as a rapid, reproducible, and low-cost alternative for species-level identification of freshwater snails (Hamlili et al., 2021). This method, capable of processing 96 samples in less than one hour, overcomes the limitations of both morphological and molecular methods (Hamlili *et al*., 2021; Gaye *et al*., 2024; Nakatt *et al*., 2024). Approximately a decade ago, MALDI-TOF MS profiling technology was applied in malacology for the identification of marine bivalves, such as scallop species (Stephan et al., 2014). This application has since been followed by very recent studies aimed at identifying certain gastropod species, notably schistosome intermediate host snails, using specimens that were either frozen, laboratory-reared, or preserved in 70% ethanol (Hamlili et al., 2021; Gaye et al., 2024; Nakatt et al., 2024). Accurate identification of medically relevant freshwater snails is a crucial initial step for establishing effective strategic control measures It has been reported that the freezing of specimens at -20°C improved the performance of MALDI-TOF MS by increasing the rate and the accuracy of snail species identification (Hamlili *et al*., 2021; Gaye *et al*., 2024). Conversely, others storing mode like ethanol offered less efficient impact for snail specimen classification based on resulting MALDI-TOF MS spectra (Hamlili *et al*., 2021; Nakatt *et al*., 2024). These same observations were reported in several arthropod genera, showing the positive effect of freezing on the generated MS spectra compared to other storing mode like ethanol or dry with silicagel at room temperature (RT) (Yssouf et al. 2014; Nebbak *et al*., 2017Briolant et al., 2020; Bamou et al., 2022; Costa et al., 2023). Although frozen specimens generate high-quality MALDI-TOF MS spectra, maintaining a cold chain under field conditions remains challenging in many endemic settings because dry ice and liquid nitrogen are often unavailable. Consequently, there is a need for preservation methods that are both operationally feasible and compatible with reliable MALDI-TOF MS identification.

The objectives of this study were therefore to: (i) evaluate the performance of MALDI-TOF MS for identifying freshwater snails preserved in 70% ethanol at 4 °C and collected from temporary and permanent water bodies in southern Mauritania; (ii) characterize the distribution of snail species across these habitats; (iii) assess infestation rates with *Schistosoma haematobium* complex parasites using real-time PCR; and (iv) optimize the pretreatment procedure for MALDI-TOF MS analysis.

## Material and methods

### Study area

The study was conducted in the villages of Kankossa and Oued Rawdha, located in the Assaba province of southern Mauritania (Supplementary Figure 1). A detailed description of these study sites was previously provided by Nakatt *et al*. (2024). Briefly, Kankossa (11,000 inhabitants) is located approximately 80 km south of Kiffa, the provincial capital, along the shores of the permanent Kankossa lake. This lake, covering approximately 7.5 km^2^, is one of the largest permanent bodies of water in Mauritania. The Oued Rawdha, a small village of 2,000 inhabitants, is situated about 8 km northwest of Kiffa, bordering a 15 km long basin of the same name. The Oued Rawdha basin contains a seasonal stream that provides water for market gardens and oasis crops during the rainy season.

### Snail collection and morphological identification

Cross-sectional malacological surveys were conducted in 2023 in the villages of Kankossa and Oued Rawdha during the rainy season (July to September). Snails were collected from the permanent lake and temporary ponds in Kankossa, and from the seasonal streams and temporary ponds in Oued Rawdha. Snail collection was conducted over two consecutive weeks between 8:00 and 10:00 am at seven sites in Kankossa and five sites in Oued Rawdha (Supplementary Table S1). Snails were collected by hand when possible or using a standard dip net. Snails from the same water point were placed in the same pre-labeled container and transported to the laboratory. All snails were morphologically identified to the species level based on shell morphology, using standard taxonomic keys (Brown & Kristensen, 1993). The snails were then sorted by species and preserved in 70% ethanol at 4°C for subsequent molecular and proteomic analyses at the Parasitology laboratory of the Institut Hospitalo-Universitaire Méditerranée Infection, Marseille, France.

### Homogenization of snails’ feet for MALDI-TOF MS analysis

To isolate the foot, the shell protecting the body of each snail was carefully removed using a sterile slide, with a separate slide used for each specimen. The dissected feet were then rinsed twice with distilled water and dried on sterile filter paper overnight at 37°C, before being homogenized according to the previously described protocol (Hamlili et al,. 2021). In 1.5 mL microcentrifuge tubes, the dissected feet pieces were placed individually, in the presence of glass beads (#11079110, BioSpec Products, Bartlesville, OK, US). A 30 µL volume of a protein extraction mix consisting of HPLC-grade water, 70% (v/v) formic acid (Sigma-Aldrich, Lyon, France) and 50% (v/v) acetonitrile (Fluka, Buchs, Switzerland), was added to each tube. The mixtures were then homogenized for three cycles of one minute each, at a frequency of 30 Hz, using a TissueLyser II (Qiagen, Hilden, Germany), after brief centrifugation of the samples for 30 seconds at 2,000 × g. On a MALDI-TOF steel target plate (Bruker Daltonics, Wissembourg, France), 1 µL of protein extract was deposited in four spots. After drying at room temperature, each spot was covered with 1 μL of a matrix solution containing of HPLC-grade water, saturated α-cyano-4 hydroxycinnamic acid (Sigma, Lyon, France), 50% (v/v) acetonitrile and trifluoroacetic acid (2.5% v/v) (Aldrich, Dorset, UK).

### MALDI-TOF MS settings and spectra acquisition

Mass spectrometry analysis was carried out using Microflex LT MALDI-TOF mass spectrometry (Bruker Daltonics machine) with FlexControl v.2.4 software (Bruker Daltonics). MALDI-TOF MS spectrometer parameters were similar to those previously described (Lafri et al., 2016). MS spectra were acquired in positive-ion linear mode over a range from 2,000 to 20,000 Da at a frequency of 50 Hz. MS spectra were generated from 240 laser shots performed at six different sites within a single spot.

### MS spectral analysis of snail feet

FlexAnalysis v.3.3 software (Bruker Daltonics) was used for visual validation of MS spectra. For spectral data processing (smoothing, base subtraction, and peak selection), plot analysis, dendrogram generation and composite correlation index (CCI) matrix construction, spectra were exported to ClinProTools v.2.2 and MALDI-Biotyper v.3.0 (Bruker Daltonics, Germany), to evaluate the reproducibility of MS spectra within the same species, as well as the specificity of MS spectra between species (for details see Costa *et al*., 2023; Almeras, *et al*., 2024). The interpretation of the CCI analysis was based on the correlation index value, which ranges from 0 to 1, and the color scale, which varies from blue to red, corresponding respectively to divergence and reproducibility of the spectra. Each MS spectra which did not reach the inclusion criteria (more intense peak > 3000 arbitrary unit (a.u.), background lower than 15-fold of the more intense peak), were considered as unconfirmed and excluded from analysis (Bouledroua *et al*., 2025).

### Database creation and blind tests

Our home-made reference MS spectra database was updated by the introduction of five representative MS spectra for each snail species (n=5). MS spectra from each specimen used as reference, identified by morphological, were validated by molecular methods. The MS reference spectra were created using an unbiased algorithm based on frequency, peak position and intensity information. MS spectra from the remaining snails not used for the database upgraded, were queried against our home-made database of reference MS spectra. Reliability of species identification was evaluated using logarithmic score values (LSVs), with a scale ranging from 0 to 3 by MALDI-Biotyper v.3.0 software (Bruker Daltonics) to ensure adequate identification, a LSV threshold upper or equal to 1.8 should be reach, as suggested previously (Briolant *et al*., 2020; Nebbak *et al*., 2021). The MALDI-TOF MS identification was considered correct when it agreed with the morphological identification and reached the LSV threshold. In case of discrepancy or inadequate identification, the specimen species was based on DNA sequences analysis.

### Snails DNA extraction and sequencing

Five specimens of each morphologically identified species —*B. truncatus, B. umbilicatus, B. forskalii* and *B. senegalensis*, and *Melanoides tuberculata —*were selected for DNA-based identification. Genomic DNA was extracted from the remaining tissue of each individual snail (excluding the foot, which was used for MALDI-TOF analysis) using the NucleoMag pathogen kit (MACHEREY-NAGEL), according to the manufacturer’s instructions using the King-Fischer apparatus. Molecular identification of snails at the species level was achieved by sequencing a 710 bp DNA fragment of the cytochrome c oxidase I (COI) gene. PCR amplification was performed using the universal primers LCO1490 (F_5’-GGTCAACAAATCATAAAGATATTGG-3’) and HC02198 (R_5’- TAAACTTCAGGGTGACCAAAAAATCA-3’), as previously described (Folmer *et al*., 1994). PCR amplification, purification of the COI gene amplicon, bidirectional Sanger sequencing (5’ to 3’ and 3’ to 5’), sequence assembly and correction using ChromasPro software version 1.7.7, BLAST analysis against the NCBI online database for species identification based on COI sequence data, phylogenetic tree construction were performed as previously described (Nakatt *et al*., 2024). The most appropriate substitution model for phylogenetic tree construction was determined using ’’Find Best DNA/Protein Models’’ option in MEGA (Molecular Evolutionary Genetics Analysis) software v7.0.26 (Tamura & Nei, 1993; Kumar *et al*., 2016). The following reference sequences, obtained from GenBank, were used in the phylogenetic analysis: *B. senegalensis*, OP811027.1 and OP811029.1; *B. umbilicatus*, OP779242.1 and OR921238.1; *B. truncatus*, MG407283.1 and MT272328.1; *B. forskalii*, MT272322.1 and ON112326.1; *B. globosus*, OR553990.1; *B. ugandae*, OP242173.1; *B. tropicus*, MN551500.1; *B. nasutus*, OP233139.1; *M. tuberculata*, MW167052.1; *M. imitatrix*, DQ995479.1; and *Lanistes varicus*, MW579648.1. Statistical support for internal branches of trees was assessed using bootstrap with 1000 replications.

### PCR detection of Schistosoma spp. infection in snails

Real-time polymerase chain reaction (RT-PCR) was used to detect and screen for S. *hæmatobium* complex parasites in the collected snails using their DNA extracts. The highly repetitive 120-base-pair DraI region, a key gene frequently used for screening this parasite complex, was targeted. The primer pairs, probe and sequences, and RT-PCR amplification conditions were in accordance with the method previously established by Nakatt *et al*. (2024). Samples with a Ct threshold of less than 35 were considered positive.

## Results

### Snail collection and morphological identification

A total of 806 live freshwater snail specimens (538 from Kankossa, 268 from Oued Rawdha) were collected from various water points during the 2023 rainy season. Based on key morphological characteristics, they were identified as *B. forskalii* (n = 102), *B. senegalensis* (n = 103), *B. truncatus* (n = 387), *B. umbilicatus* (n = 165) and *Melanoides tuberculata* (n = 49) (Table 1). Although several snail species were found per locality, none of the five species was detected at both study sites.

**Table 1.** Morphological identification of freshwater snails collected during the 2023 rainy season at Kankossa and Oued Rawdha.

| Species | Number of specimens collected per study site (%) |  |
| --- | --- | --- |
|  | Kankossa | Oued Rawdha |
| *Bulinus forskalii* | 102 (12.7) | NF |
| *Bulinus senegalensis* | NF | 103 (12.8) |
| *Bulinus truncatus* | 387 (48.0) | NF |
| *Bulinus umbilicatus* | NF | 165 (20.5) |
| *Melanoides tuberculata* | 49 (6.1) | NF |
| **Total** | **538 (66.8)** | **268 (33.3)** |

NF: Not found.

### DNA sequence-based identification of selected snails

A subgroup of 25 snails, five per species based on morphological identification were submitted to Cytochrome Oxidase I (COI, 710 bp) gene sequencing to validate their classification. Sequence analysis results confirmed morphological results (Table 2). BLAST analysis of COI sequences yielded high identity scores ranging from 98.9% to 100% (Table 2). The phylogenetic tree based on COI sequences demonstrated clear discrimination between the five species (Figure 1). Furthermore, each species formed a distinct group that associated with homologous sequences available in GenBank on the same phylogenetic branch.

**Figure 1.**
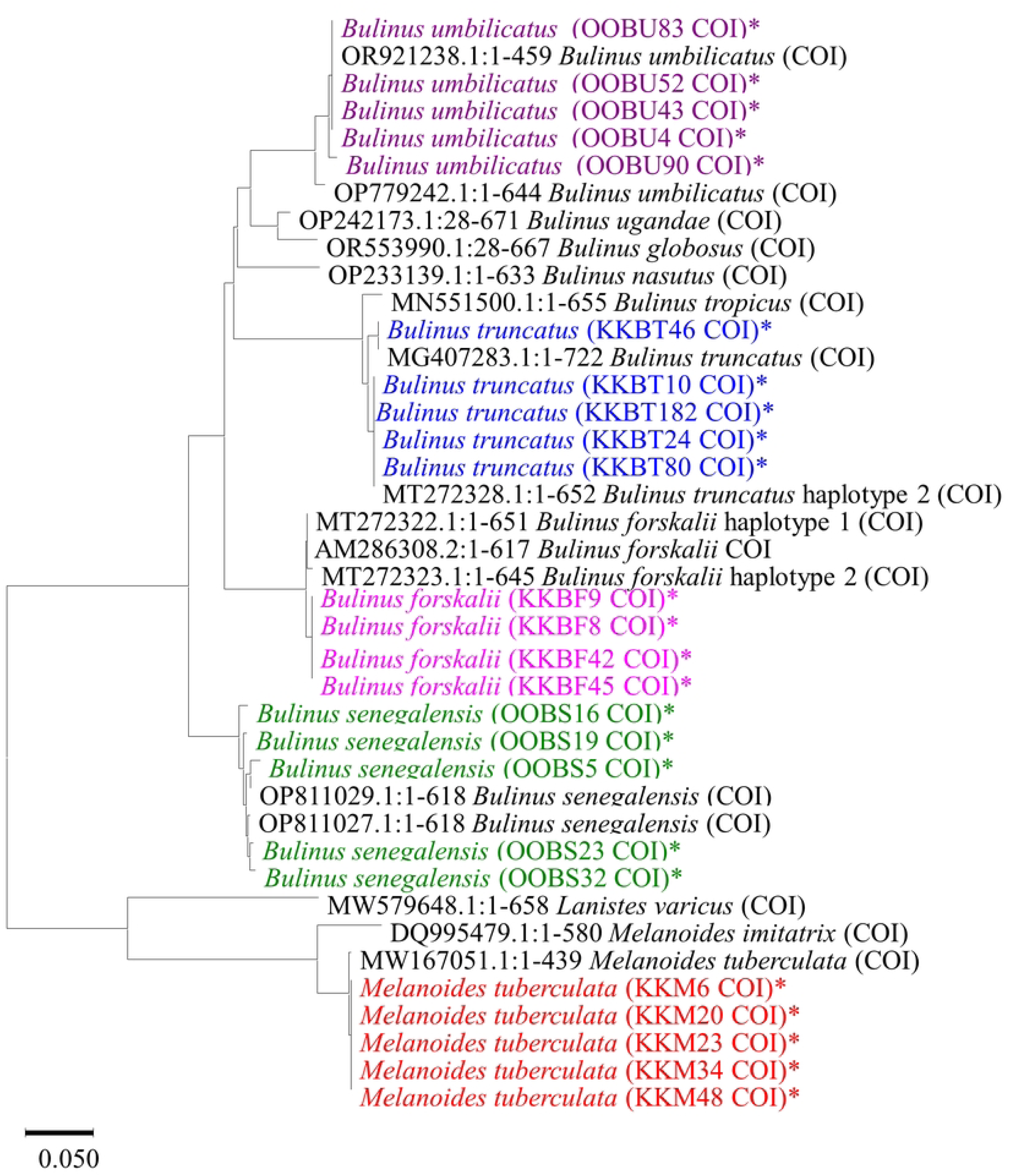
Phylogenetic tree based on the nucleotide sequence of a 710-base pair region of the mtDNA. The phylogenetic tree was constructed using maximum likelihood method based on the TN93+G substitution model with MEGA 7. The values on the branches are bootstrap support values based on 1000 replicates. The asterisks indicate the specimens collected and analyzed in the present study with their corresponding GenBank accession numbers. The other snails are reference specimens with their GenBank accession number.

**Table 2.** Identification of *Bulinus* spp. and *Melanoides tuberculata* snail species based on mitochondrial DNA (COI) sequences.

| Morphological ID | Number of specimens tested per species | COI Mitochondrial Genome |  |  |
| --- | --- | --- | --- | --- |
|  |  | Species ID | Range of identity (%)<sup>(a)</sup> | NCBI Acc. No<sup>(b)</sup> |
| *B. forskalii* | 5 | *B. forskalii* | 99.4 - 99.7 | MT272322.1,<br>AM286308.2 |
| *B. senegalensis* | 5 | *B. senegalensis* | 98.9 - 99.7 | OP811027.1,<br>OP811028.1,<br>OP811029.1 |
| *B. truncatus* | 5 | *B. truncatus* | 99.5 – 100 | MG407283.1,<br>MG407261.1,<br>MG407285.1,<br>KJ135308.1 |
| *B. umbilicatus* | 5 | *B. umbilicatus* | 98.9 – 100 | OP779242.1,<br>OR921238.1 |
| *M. tuberculata* | 5 | *M. tuberculata* | 99.8 | MW167051.1 |

ID: Morphological Identification; n: number of snail specimens that have been sequenced. (a): best sequence identity score; (b): NCBI GenBank accession numbers of nucleotide sequences with the highest identity percentage.

### Assessment of snail MS spectra reproducibility and specificity per species

Prior to evaluate the MALDI-TOF MS classification performances, the MS spectra from the 25 specimens validated by gene sequencing (ie, *B. senegalensis* (n = 5), *B. truncatus* (n = 5), *B. umbilicatus* (n = 5), *B. forskalii* (n = 5) and *M. tuberculata* (n = 5), Table 2), were compared. The visual comparison of these 25 MS spectra showed a high intra-species reproducibility and inter-species specificity for the four *Bulinus* species (Figure 2A). Interestingly, despite an overall reproducibility of *M. tuberculata* MS spectra, several intense supplementary peaks at above 11000 Da, notably at 11355 Da and 13580 Da, were detected in two out of five specimens (Figure 2A).

**Figure 2.**
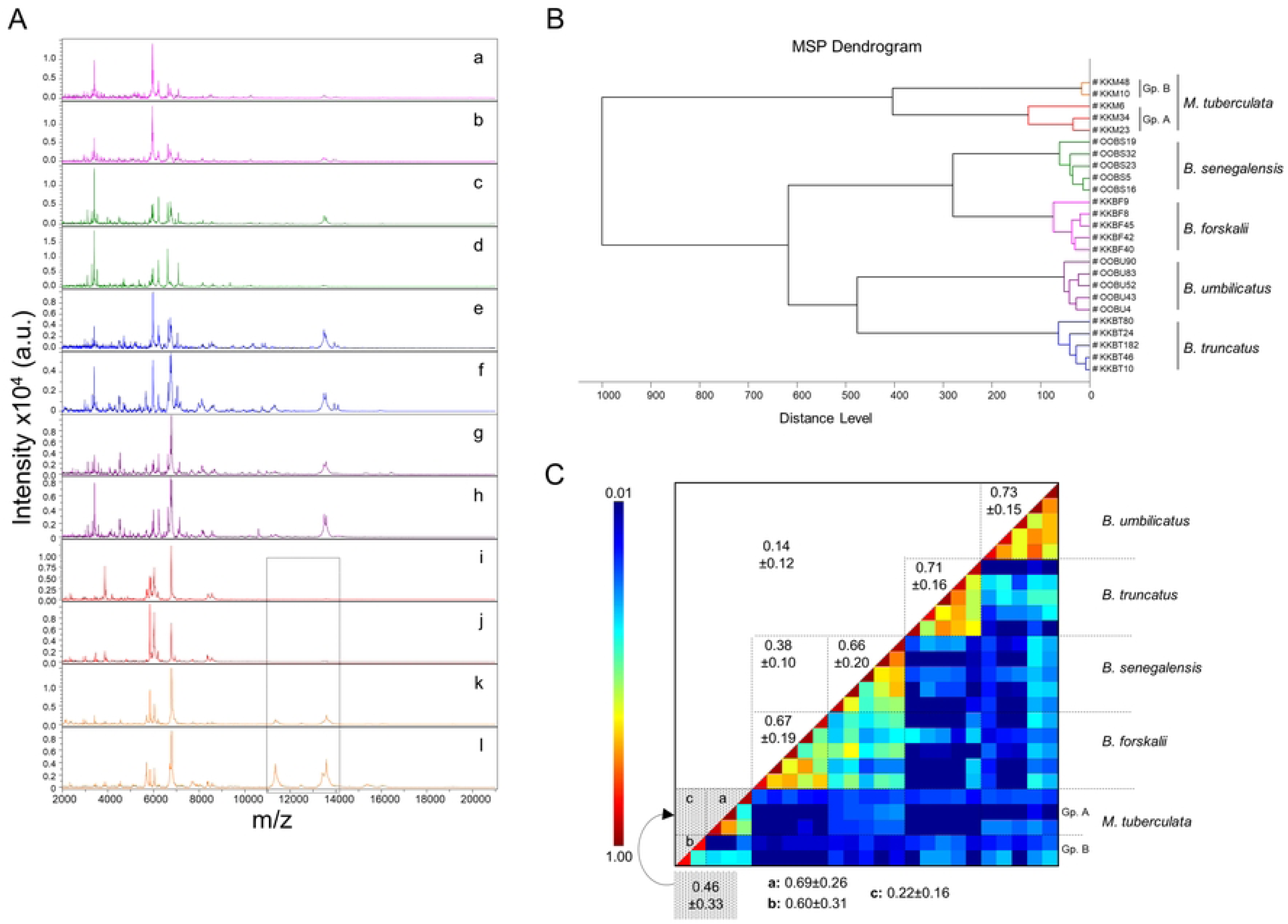
MALDI-TOF MS spectra from five freshwater snail species. (**A**) Representative MS spectra from *Bulinus* spp. (*B. forskalii* (a, b), *B. senegalensis* (c, d), *B. truncatus* (e, f), *B. umbilicatus* (g, h), and *Melanoides tuberculata* (i, j, k, l). Divergent intra-species MS peaks are surrounded by a black frame. (**B**) MSP dendrogram of MALDI-TOF MS spectra of snail, using 25 selected *Bulinus* spp. and *M. tuberculata* specimens as reference spectra. The distance units correspond to the relative similarity of MS spectra. The dendrogram was created by Biotyper v3.0 software. (**C**) The same 25 MS spectra were analyzed using the composite correlation index (CCI) tool. CCI matrix illustrates the homogeneity and reproducibility levels of MS spectra within the same species and between different species. Reproducibility levels of the MS spectra are indicated in red and blue, revealing relatedness and incongruence between spectra, respectively. Values correspond to the mean correlation coefficient and respective standard deviations obtained for paired condition comparisons and were expressed as mean ± standard deviation. CCI was calculated using MALDI-Biotyper v.3.0 software. In panel (B) and (C), spectra from *M. tuberculata* specimens were clustered into two subgroups “Gp. A” and “Gp. B” according to their MS spectra. *B*., *Bulinus*; *M*., *Melanoides*; a.u., arbitrary units; MSP, Main Spectrum Profile; m/z, mass-to-charge ratio; SD, standard deviation; GP., group.

To assess whether the species clustered according to their MALDI-TOF spectra, a MSP dendrogram and CCI matrix were done. The MSP dendrogram revealed that the primary clustering occurred at the genus level, separating *Bulinus* spp. from *Melanoides tuberculata* (Figure 2B). It also disclosed that MS spectra from specimens of the same species were grouped on the same branch, confirming the species homogeneity of MS spectra and their distinction among species (Figure 2B). The high CCI values obtained for *Bulinus* specimens of the same species, ranging from 0.66 ± 0.24 (mean ± SD) for *B. senegalensis* to 0.73 ± 0.15 for *B. umbilicatus*, supporting the intra-species homogeneity of MS spectra (Figure 2C). Among the comparisons of paired MS spectra between *Bulinus* species, the higher correlation (mean ± SD: 0.38 ± 0.10) was obtained between *B. forskalii* and *B. senegalensis*. However, this CCI was low compared to those from intra-species comparisons preventing the risk of future misidentification.

For spectra from *M. tuberculata* specimens, a low CCI (mean ± SD: 0.46 ± 0.29) was recovered compared to *Bulinus* species. This lower CCI value was attributed to the heterogeneity of spectra among the *M. tuberculata* specimens, as noticed on MS spectra (Figure 2A). The MSP dendrogram separated *M. tuberculata* spectra into two subgroups, arbitrary named “Group B” and “Group A”, accordingly to the presence or not of intense peaks above 11000 Da, respectively. The application of this clustering on CCI analysis revealed higher intra-group values (ie, mean ± SD of 0.69 ± 0.26 and 0.60 ± 0.31 for Groups A and B, respectively). Whereas, very low values (ie, mean CCI < 0.23) for paired MS spectra comparison between groups A and B were obtained. These results underlined that snails from M*. tubercularta* species could be separated into two subgroups based on MALDI-TOF MS analysis of resulting spectra. Each subgroup presenting homogeneous spectra and sufficiently specific to generate low inter-group correlation value. The origin of this intra-species heterogeneity remains at this stage unknown.

### Database creation and blind tests

The spectra from the 25 specimens from the five snail species, validated by COI sequencing and producing MS spectra of high intensity, were selected for creation of reference MS spectra. These MS spectra were added to our homemade MS spectra database using MALDI-Biotyper 3.0. (Additional file 1). The remaining snail feet (n=781) were submitted to MALDI-TOF MS analysis. Among them, MS spectra of high-intensity were obtained for 99.0% (773/781) of the samples tested. The MS spectra from these eight specimens, morphologically identified as *B. truncatus* (n=3), *B. umbilicatus* (n=3), and *M. tuberculata* (n=2) which did not reach the inclusion criteria (i.e., most intense peak lower than 3000 a.u. and high background) were considered as irrelevant and were excluded from the analysis.

The spectra from the remaining specimens (n=773) were blind tested against our updated MS database (Table 3). Overall, the proportion of concordant identification by MALDI-TOF MS with morphological classification was 99.1% (766/773). Among the specimens concordantly identified, nearly 97.0% (743/766) reached the LSVs > 1.8 threshold value to consider the identification as adequate (Figure 3). *B. senegalensis* was the unique species for which identification at the species level were 100% concordant with morphology and LSVs>1.8. Interestingly, for the four *Bulinus* species, the mean LSVs were all above 2.3 highlighting the high specificity of the MS spectra and confidence in species identification. Finally, a total of 30 samples, representing less than 3.9% of the spectra, either failed to reach adequate threshold value (n=23), or conducted to discrepant identification (n=6), or both (n=1).

**Figure 3.**
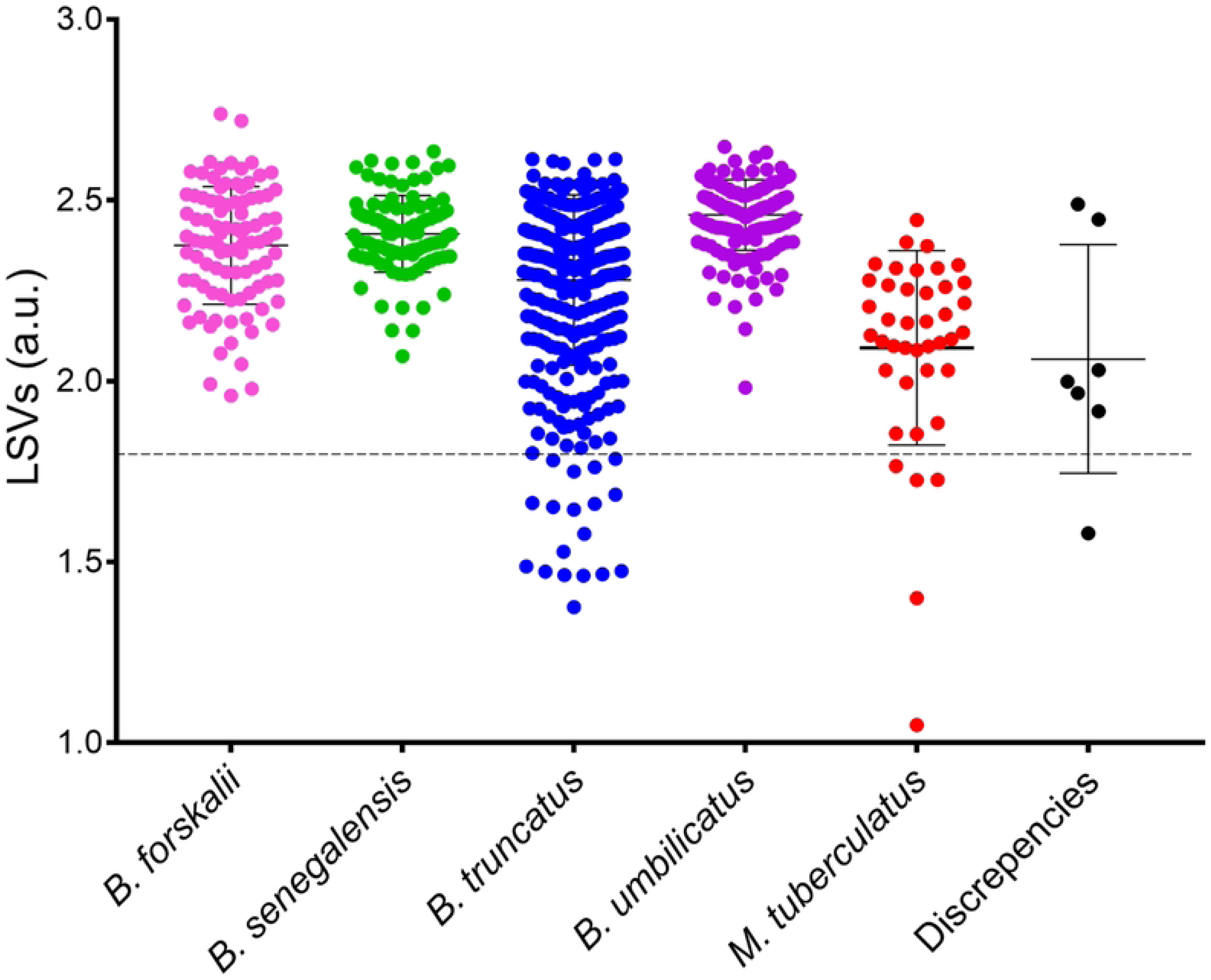
LSVs of the 773 snails MS spectra query against the updated MS reference database. Each dot represent an identification result per species with it respective LSV. Discrepant identification results between MS and morphological classification are indicated by black dots. LSVs were expressed as mean ± standard deviation for species of *Bulinus spp.* And *M. tuberculata*. The horizontal dashed line represents the threshold value (≥ 1.8) for adequate identification. a.u., arbitrary units; LSV, log score value.

**Table 3.** MALDI-TOF MS-based identification of the *Bulinus* and *Melanoides* snail species from the lake zones of Kankossa and Oued Rawdha, in the Wilaya of Assaba, southern Mauritania.

| **Morphological ID (n)** | **MS spectra of good quality (%)** | **MS spectra used as reference (n)\*** | **Spectra blind tested (n)** | **MS ID (n)** | **Range of LSVs** | **Adequate and concordant ID (%)#** |
| --- | --- | --- | --- | --- | --- | --- |
| *B. forskalii* (102) | 102 (100) | 5 | 97 | *B. forskalii* (95) | [1.96-2.74] | 95 (100) |
|  |  |  |  | *B. truncatus* (2) | [1.98-2.03] | 2 (100) |
| *B. senegalensis* (103) | 103 (100) | 5 | 98 | *B. senegalensis* (98) | [2.07-2.64] | 98 (100) |
| *B. truncatus* (387) | 384 (100) | 5 | 379 | *B. truncatus* (377) | [1.38-2.61] | 359 (95.2) |
|  |  |  |  | *B. forskalii* (2) | [2.45-2.49] | 2 (100) |
| *B. umbilicatus* (165) | 162 (98.8) | 5 | 157 | *B. umbilicatus* (155) | [1.92-2.65] | 155 (100) |
|  |  |  |  | *B. senegalensis* (2) | [1.92-1.97] | 2 (100) |
| *M. tuberculata* (49) | 47 (95.9) | 5 | 42 | *M. tuberculata* (41) | [1.05-2.45] | 36 (87.8) |
|  |  |  |  | *B. truncatus* (1) | [1.58] | 0 (0.0) |
| **Total (806)** | **798** | **25** | **773** |  |  | **749 (97.1)** |

\*The species identity of all specimens included in the reference MS spectra DB were validated by DNA sequencing; #Are considered as adequate if MS spectra LSV was greater than 1.8 and concordant with morphological ID; ID, identification; n, number; MS, mass spectrometry; LSVs, log score values.

### Molecular identification of discrepant and/or inadequate results

For the thirty snails for which discrepant and/or inadequate results were obtained, the COI gene was sequenced to determine species identification (Table 4). COI gene sequencing was successful for 29 out of 30 snails tested. The failure of molecular identification occurred for one *M. tuberculata* sample for which MS identification was discordant and inadequate. For the 29 remaining samples, DNA sequencing confirmed the MS classification for the six discordant identification with morphology and for 23 inadequate MS classification. Then, overall, COI gene sequencing revealed that MALDI-TOF MS identified successfully 99.9% (n=772/773) of the snails, corroborating either morphological or molecular results in case of discrepancies.

**Table 4.** Snail specimens submitted for molecular identification using the COI gene sequence to complete Mass Spectrometry identification.

| **Morphological ID (n)** | **Reason for molecular identification<sup>#</sup> (n)** | **MS ID (n)** | **Molecular ID (n)** | **Range of identity (%)<sup>(a)</sup>** | **NCBI Acc. No<sup>(b)</sup>** |
| --- | --- | --- | --- | --- | --- |
| *B. forskalii* (2) | Discordant (2) | *B. truncatus* (2) | *B. truncatus* (2) | 99.4 - 99.7 | MT272328.1,<br>MT272331.1 |
| *B. truncatus* (20) | Inadequate (18) | *B. truncatus* (18) | *B. truncatus* (18) | 99.8 - 100 | MT272328.1,<br>MG407285.1,<br>KR108924.1 |
|  | Discordant (2) | *B. forskalii* (2) | *B. forskalii* (2) | 99.5- 99.7 | AM286308.2 |
| *B. umbilicatus* (2) | Discordant (2) | *B. senegalensis* (2) | *B. senegalensis* (2) | 99.5 | OP811028.1,<br>MW167056.1 |
| *M. tuberculata* (6) | Inadequate (5) | *M. tuberculata* (5) | *M. tuberculata* (5) | 99.5 - 99.8 | MW167051.1 |
|  | Disc. & Irr. (1) | *B. truncatus* (1) | NI |  |  |
| Total | 30 |  | 29 |  |  |

<sup>#</sup>Three reasons were selected: *Discordant* corresponding to discordant identification between morphology and MALDI-TOF MS; *Inadequate* corresponding to Log Score Value below the threshold of 1.8 to consider ID as adequate. ID: Identification; MS: mass spectrometry; No: number; NI: Not identified.

### Molecular screening of snails infested with *S. hæmatobium* complex parasites

A molecular screening of the *S. haematobium* complex was performed on 798 snail specimens exhibiting good quality MS spectra. The results revealed the presence of the *S. haematobium* complex in 239 (29.9%) of these specimens (Table 5). Among them, 16 specimens came from Oued Rawdha, (9 *B. senegalensis* and 7 *B. umbilicatus*) and 223 specimens came from Kankossa (173 *B. truncatus*, 34 *B. forskalii* and 16 *M. tuberculata*). The infestation rates of the *S. haematobium* complex were higher in permanent watercourses of Kankossa (41.8%, n=223/533) compared the temporary pool of Oued Rawdha (6.0%, n=16/265). Cycle threshold (Ct) values ranged from 4.12 to 34.46. The Ct values indicated that the *Melanoides* snails had relatively low parasitic loads compared with the *Bulinus* snail (Figure 4).

**Figure 4.**
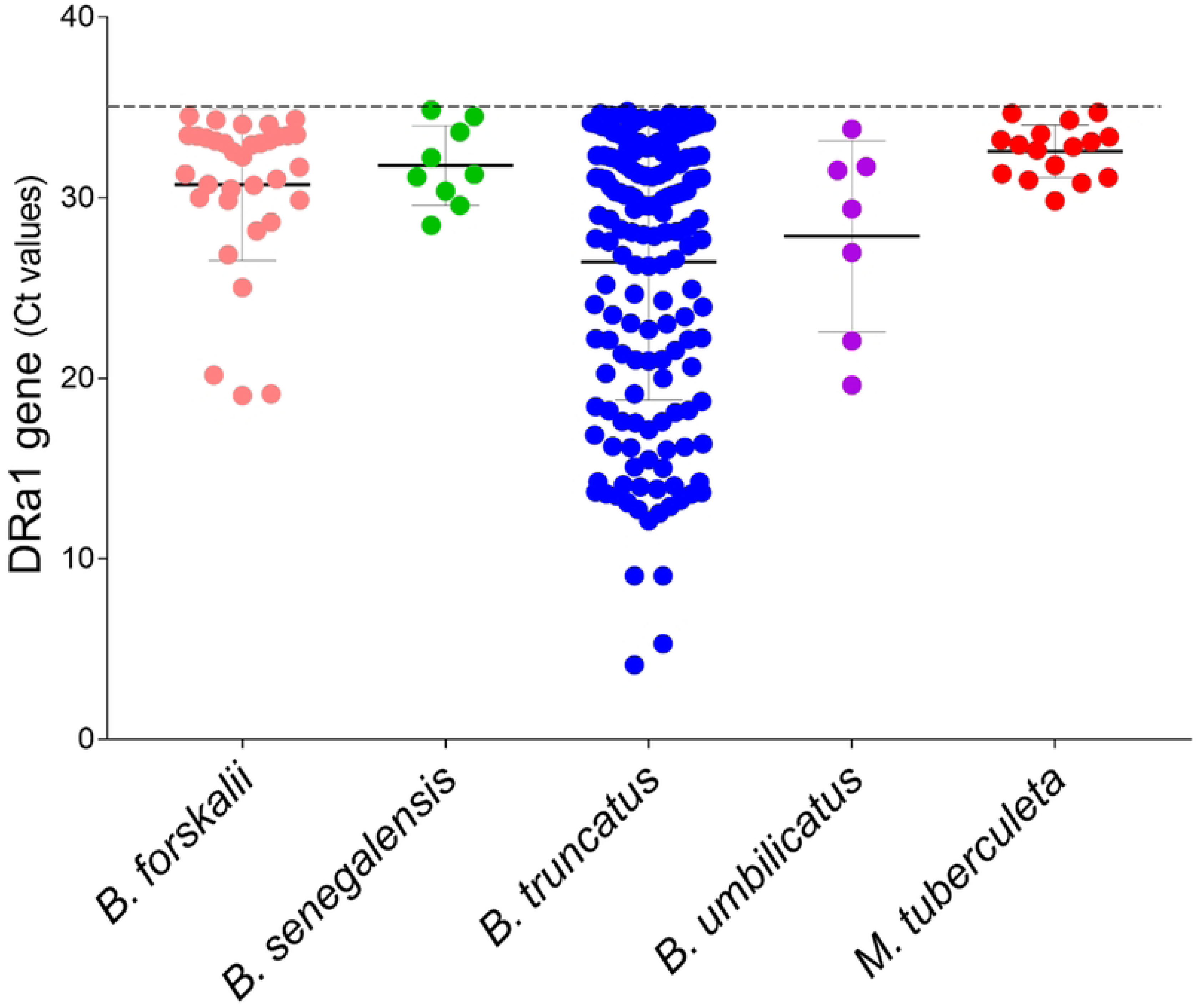
Cycle threshold values *Bulinus* spp. and *M. tuberculata* infected by *Schistosoma haematobium* complex using qPCR detection of the Dra1 region. Ct values were expressed as mean ± standard deviation for species of *Bulinus* spp. and *M. tuberculata*. The dashed line represents the threshold value (< 35) for considering the sample positive.

**Table 5.** *Schistosoma haematobium* complex infestation rate in *Bulinus* spp. and *M. tuberculata* using qPCR detection of the Dra1 region.

| Snail species | Sample tested (n) | *S. haematobium* infection (n) | Proportion (%) |
| --- | --- | --- | --- |
| *B. forskalii* | 102 | 34 | 33.3 |
| *B. senegalensis* | 105 | 9 | 8.6 |
| *B. truncatus* | 384 | 173 | 45.1 |
| *B. umbilicatus* | 160 | 7 | 4.4 |
| *M. tuberculata* | 47 | 16 | 34.0 |
| Total | 798 | 239 | 29.9 |

*B.*, *Bulinus* ; *M.*, *Melanoides*; *S.*, *Schistosoma*

### Assessment of snail *Schistosoma* spp. infection using MALDI-TOF MS

To determine whether MALDI-TOF MS could distinguish *Schistosoma-*infected from non-infected snails, a comparison of MS spectra were done. Paired comparison per species using ClinProTools v.2.2 (Bruker Daltonics) did not allowed to visualize any MS peaks in the *Schistosoma* infected group versus non-infected snails of the same species. Moreover, PCAs done using these spectra datasets, supported the absence of MS peak(s) separating snails according to their infectious status (Supplementary Figure S2A-E). It is interesting to note that MS peaks above 11 kDa found in a subgroup of *M. tuberculata* specimens (Group B) were found in both *Schistosoma-*infected and non-infected snails (Supplementary Figure S2F). These peaks could then likely considered as independent from *Schistosoma* infection of *M. tuberculata* specimens.

## Discussion

The present study demonstrates that MALDI-TOF MS can accurately identify medically relevant freshwater snails collected under field conditions and preserved in 70% ethanol at 4 °C prior to analysis. This optimized preservation strategy yielded high-quality spectra for nearly all specimens analyzed and markedly improved identification performance compared with previous studies using ethanol preservation at room temperature. These findings are operationally important for schistosomiasis-endemic regions where maintaining frozen storage conditions during field surveys is often impractical. Previous studies reported that frozen specimens generated the most reproducible and informative MALDI-TOF MS spectra for freshwater snail identification (Hamlili *et al*., 2021; Gaye *et al*., 2024). However, access to dry ice, liquid nitrogen, or continuous cold-chain infrastructure remains limited in many endemic settings. In the present work, storage in 70% ethanol at 4 °C enabled successful acquisition of high-intensity spectra for 99.0% of the analyzed specimens, whereas only eight samples failed to meet predefined quality criteria. This performance is substantially higher than previously reported for ethanol-preserved snails stored at room temperature (Hamlili *et al*., 2021; Nakatt *et al*., 2024), and approaches the performance reported for frozen or laboratory-reared specimens. The improved results obtained with storage at 4 °C likely reflect better preservation of protein integrity during storage.

MALDI-TOF MS identification showed excellent agreement with morphological and molecular classification. After molecular confirmation of discrepant and low-confidence results, the overall identification accuracy reached 99.9%. These findings highlight the robustness of MALDI-TOF MS for freshwater snail identification and support its application as an alternative to conventional morphology-based approaches. Morphological identification of freshwater snails may be challenging in field conditions because shell characteristics can vary according to environmental conditions, developmental stage, or shell erosion. In particular, species belonging to the *Bulinus forskalii* complex, especially *B. forskalii* and *B. senegalensis*, are notoriously difficult to distinguish using shell morphology alone (Jobin *et al*., 1976; Betterton *et al*., 1983). In the present study, no misclassification was observed between these two species using MALDI-TOF MS, further supporting the discriminatory power of this proteomic approach. Compared with molecular methods, MALDI-TOF MS also presents several operational advantages. DNA-based identification remains reliable but requires relatively expensive reagents, longer processing times, and specialized laboratory infrastructure. In addition, publicly available sequence databases for freshwater snails remain incomplete for several taxa. Once a reference database is established, MALDI-TOF MS enables rapid, high-throughput, and cost-effective species identification, making it particularly suitable for large-scale epidemiological surveillance.

Distinct ecological distributions of snail species were observed between the two study areas. *Bulinus senegalensis* and *B. umbilicatus* were exclusively detected in temporary ponds in Oued Rawdha, whereas *B. truncatus*, *B. forskalii*, and *Melanoides tuberculata* were found in the permanent water bodies of Kankossa. These findings are consistent with previous ecological observations in West Africa indicating that *B. senegalensis* and *B. umbilicatus* are well adapted to seasonal habitats and are capable of surviving prolonged dry periods through aestivation (Greer *et al*., 1990; Senghor *et al*., 2015; Senghor *et al*., 2023). In contrast, *B. truncatus* is more commonly associated with perennial water bodies (Sène *et al*., 2004; Hamlili *et al*., 2021). Such ecological differences are likely influenced by multiple environmental factors, including hydrological stability, physicochemical water composition, vegetation, and interspecific competition. Additional ecological investigations are required to better characterize the determinants of snail distribution and their impact on schistosomiasis transmission.

Interestingly, MALDI-TOF MS analysis revealed heterogeneity among *M. tuberculata* spectra, with two reproducible spectral subgroups distinguished by the presence or absence of intense peaks above 11 kDa. These spectral differences were independent of *Schistosoma* infection status and may reflect underlying biological or environmental variability within the species. The origin of this heterogeneity remains unclear and warrants further ecological and genomic investigation.

qPCR screening revealed relatively high levels of *Schistosoma haematobium* complex infection among snails collected in Kankossa compared with Oued Rawdha. The markedly higher infestation rates observed in the permanent lake of Kankossa may reflect sustained contamination of water bodies and continuous exposure of snails to parasite transmission cycles throughout the year. Conversely, the lower prevalence observed in temporary ponds at Oued Rawdha may be related to the seasonal and unstable nature of these habitats. Similar infestation patterns have previously been reported in Mauritania and neighboring West African countries (Ouldabdallahi *et al*., 2010; Tian-Bi *et al*., 2019).

Although MALDI-TOF MS has successfully been used to detect pathogen-associated spectral modifications in several arthropod vectors, the present study did not identify reproducible spectral differences between infected and non-infected snails. Principal component analyses failed to discriminate specimens according to their *Schistosoma* infection status within each species. Several factors may explain this limitation, including the relatively low parasite biomass compared with host proteins and the heterogeneity of infection intensity among specimens. The relatively high Ct values observed in many positive samples further suggest low parasite loads in several infected snails. Additional optimization strategies may therefore be necessary before MALDI-TOF MS can reliably detect *Schistosoma* infection directly from snail protein spectra.

The detection of *S. haematobium* complex DNA in *M. tuberculata* specimens should be interpreted cautiously. Detection of parasite DNA alone does not demonstrate that this species acts as a competent intermediate host capable of supporting complete intramolluscan parasite development. The detected DNA may instead reflect transient exposure or non-viable parasite stages. Additional experimental and parasitological studies are therefore required to clarify the epidemiological significance of this observation.

This study has several limitations. First, the optimal duration of protein preservation in 70% ethanol at 4 °C was not specifically investigated. Nevertheless, we assessed that reliable MALDI-TOF MS identification remained possible after up to five months of storage. Second, the ecological characterization of water bodies remained limited and did not allow detailed analysis of environmental determinants associated with snail distribution. Finally, the biological significance of the spectral heterogeneity observed within *M. tuberculata* could not be resolved within the scope of the present work.

Overall, the present findings support the use of MALDI-TOF MS as a reliable and field-compatible tool for freshwater snail identification in schistosomiasis-endemic areas. The combination of ethanol preservation at 4 °C with MALDI-TOF MS analysis provides an operationally feasible strategy for large-scale malacological surveillance in resource-limited settings. Such approaches could substantially improve monitoring of intermediate host populations and contribute to integrated schistosomiasis control programs.

## Conclusion

Accurate identification of freshwater snails is essential for understanding schistosomiasis transmission dynamics and improving epidemiological surveillance in endemic areas. This study demonstrates that MALDI-TOF MS enables rapid and reliable identification of medically important freshwater snails preserved in 70% ethanol at 4 °C under field-compatible conditions. The method accurately discriminated closely related species, including *B. forskalii* and *B. senegalensis*, which are often difficult to distinguish morphologically.

Distinct ecological distributions were observed between temporary and permanent water bodies, and molecular screening revealed substantial levels of *Schistosoma haematobium* complex infection among several snail species. Overall, these findings support the use of MALDI-TOF MS as a promising tool for large-scale malacological surveillance and monitoring of schistosomiasis transmission in endemic settings.

## Declarations

### Ethics approval and consent to participate

This study was approved by the Ethics Committee of the Mauritanian Ministry of Health and reviewed by the Department for the Fight Against Neglected Tropical Diseases and Blindness, number: 059/MS/DGS/DLCMT/SLMTNC. Date: 03/10/2021.

This study did not involve any human participant or vertebrate animal requiring specific ethical approval or informed consent.

### Availability of data and materials

The raw MALDI-TOF MS reference spectra included in the database of this study are freely accessible in the Additional File 1.

### Authors’ contributions

Conceived and designed the experiments: LN, LA and SR. Analyzed the data: LN, LA, AZD, and SR. Contributed reagents/materials/analysis tools: LN, OMM and AOMSB. Drafted the paper: LN, LA and SR. Revised critically the paper: all authors.

## Acknowledgments

We would like to thank the Fondation Méditerranée Infection (FMI), which offered a personnel grant to LN.

---

**Supplementary Figure S1.** Map of the study sites (red dots) in the lake zones of Kankossa and Oued Rawdha, Assaba region, southern Mauritania. Kiffa is the regional capital of Assaba. The base layer for the map was sourced from: https://d-maps.com/carte.php?num_car=26766&lang=en.

**Supplementary Figure S2. Paired comparison per species of MALDI-TOF MS spectra between infected and no infected snails by *S. haematobium* complex.** Assessment of MS spectra variations per snail species according to infectious status was assessed using principal component analysis (PCA). PCA results from *B. forskalii* (**A**), *B. truncatus* (**B**), *B. senegalensis* (**C**), *B. umbilicatus* (**D**), and *M. tuberculata* (**E**) infected (red dots) or not infected (green dots) by *S. haematobium* complex are presented. (**F**) Gel view of *M. tuberculata* MS spectra grouped according to their infectious status. MS spectra of snails not infected (i) and infected by *S. haematobium* complex (ii) were compared. Divergent intra-species MS peaks are surrounded by a black frame. Spectra number (Sp.#) is indicated at left and peak intensity is illustrated by a grey scale in arbitrary units.

## Supplementary data

**Supplementary table S1.** GPS coordinates of sample sites.

| Location | Collection name | Latitude | Longitude |
| --- | --- | --- | --- |
| Kankossa | Site 1 | 15.938676 | -11.520716 |
|  | Site 2 | 15.939280 | -11.519396 |
|  | Site 3 | 15.940163 | -11.517495 |
|  | Site 4 | 15.940745 | -11.516534 |
|  | Site 5 | 15.937738 | -11.526208 |
|  | Site 6 | 15.938674 | -11.530165 |
|  | Site 7 | 15.938725 | -11.529121 |
| Oued Rawdha | Site 8 | 16.637432 | -11.430952 |
|  | Site 9 | 16.684221 | -11.487303 |
|  | Site 10 | 16.688835 | -11.509942 |
|  | Site 11 | 16.679212 | -11.463943 |
|  | Site 12 | 16.676197 | -11.454508 |

## References

1. Abe EM, Guan W, Guo YH, Kassegne K, Qin ZQ, Xu J, et al. Differentiating snail intermediate hosts of *Schistosoma* spp. using molecular approaches: fundamental to successful integrated control mechanism in Africa. Infect Dis Poverty. 2018;7(1):29.

2. Albarran-Melze NC, Rangel-Ruiz LJ, Gamboa-Aguilar J. Distribution and abundance of *Melanoides tuberculata* (Gastropoda: thiaridae) in the Biosphere reserve of Pantanos de Centla, Tabasco, Mexico. Acta Zoológica Mexicana. 2009;25(1):93–104. doi: 10.21829/azm.2009.251599.

3. Almeras L, Costa MM, Amalvict R, Guilliet J, Dusfour I, David JP, Corbel V. Potential of MALDI-TOF MS biotyping to detect deltamethrin resistance in the dengue vector Aedes aegypti. PLoS One. 2024;19(5):e0303027. doi: 10.1371/journal.pone.0303027. PMID: 38728353; PMCID: PMC11086877.

4. Anyan WK, Abonie SD, Aboagye-Antwi F, Tettey MD, Nartey LK, Hanington PC, Anang AK, Muench SB. Concurrent *Schistosoma mansoni* and *Schistosoma hæmatobium* infections in a peri-urban community along the Weija dam in Ghana: A wake up call for effective National Control Programme. Acta Trop. 2019;199:105116. doi: 10.1016/j.actatropica.2019.105116.

5. Ayanda OI. Prevalence of snail vectors of schistosomiasis and their infection rates in two localities within Ahmadu Bello University (A.B.U.) Campus, Zaria, Kaduna State, Nigeria. Journal of Cell and Animal Biology. 2009;3 (4): 058–061.

6. Babangida SM, Rukaiya HA, Abubakar BS, Muhammad IA. Prevalence of snail vectors of schistosomiasis along Antau River, Keffi, Nasarawa State. International journal of nature and science advance res. 2024;06(9):224–234.

7. Bamou R, Costa MM, Diarra AZ, Martins AJ, Parola P1, Almeras L. Enhanced procedures for mosquito identification by MALDI-TOF MS. Parasites & Vectors. 2022;15(240):17. 10.1186/s13071-022-05361-0

8. Banyigyi AH, Isah MH, Ameh SM, Ibrahim JO. A Survey of Freshwater Snails in Three Rivers of Keffi Local Government, Nasarawa State, Nigeria. Savanna Journal of Basic and Applied Sciences. 2022;5(1):66–71.

9. Barroso CX, Matthews-Cascon H. Occurrence of the exotic freshwater snail *Melanoides tuberculatus* (Mollusca: Gastropoda: Thiaridae) in an estuary of north-eastern Brazil. Mar. Biodivers. Rec. 2009;2,e116. 10.1017/S175526720900097

10. Barros MRF, Chagas RA, Herrmann M, Bezerra AM. New record of the invasive snail *Melanoides tuberculata* (Gastropoda, Thiaridae) - Ceará State, Brazil. Braz. J. Biol. 2020;80: 368–372. 10.1590/1519-6984.210408

11. Benyahia H, Ouarti B, Diarra AZ, Boucheikhchoukh M, Meguini MN, Behidji M, Benakhla A, Parola P, Almeras L. Identification of Lice Stored in Alcohol Using MALDI-TOF MS. J Med Entomol. 2021;58(3):1126–1133. doi: 10.1093/jme/tjaa266. PMID: 33346344.

12. Bertrand A. Mollusques terrestres et d’eau douce des Pyrénées-Atlantiques : catalogue commenté des espèces, espèces patrimoniales, enjeux de connaissances et de conservation, bibliographie. Folia Conchyliologica. 2020;55 :77.

13. Betterton C, Fryer SE, Wright CA. *Bulinus senegalensis* (Mollusca: Planorbidae) in northern Nigeria. Ann Trop Med Parasitol. 1983;77(2):143–9. doi: 10.1080/00034983.1983.11811689. PMID: 6882063.

14. Bogéa T, Cordeiro FM, Gouveia JS. *Melanoides tuberculatus* (Gastropoda: Thiaridae) as intermediate host of Heterophyidae (Trematoda: Digenea) in Rio de Janeiro metropolitan area, Brazil. Rev Inst Med Trop Sao Paulo. 2005;47(2):87–90. doi: 10.1590/s0036-46652005000200005.

15. Briolant S, Costa MM, Nguyen C, Dusfour I, Pommier de Santi V, Girod R, et al. Identification of French Guiana anopheline mosquitoes by MALDI TOF MS profiling using protein signatures from two body parts. PloS ONE. 2020;15:e0234098.

16. Brown DS, Kristensen TK. A field guide to African freshwater snails. 1. West African species. Danish Bilharziasis Laboratory: Charlottenlund, Denmark, 1993. Available online: https://www.eliminateschisto.org/sites/gsa/files/content/attachments/2022-08-05/FG_1993_West_Africa.pdf. Accessed on 13 January 2025.

17. Brown DS. Freshwater Snails of Africa and their Medical Importance. Revised 2^nd^ edition. Taylor & Francis Ltd., London, UK. 1994. Available online : https://archive.org/details/freshwatersnails0000brow/mode/2up (Accessed on 13 January 2025).

18. Bouledroua R, Diarra AZ, Amalvict R, Berenger JM, Benakhla A, Parola P, Almeras L. Assessment of MALDI-TOF MS for Arthropod Identification Based on Exuviae Spectra Analysis. Biol Proced Online. 2025;27(1):12. doi: 10.1186/s12575-024-00260-3. PMID: 40186096; PMCID: PMC11971817.

19. Costa MM, Guidez A, Briolant S, Talaga S, Issaly J, Naroua H, Carinci R, Gaborit P, Lavergne A, Dusfour I, Duchemin JB, Almeras L. Identification of Neotropical *Culex* Mosquitoes by MALDI-TOF MS Profiling. Trop Med Infect Dis. 2023;8(3):168. doi: 10.3390/tropicalmed8030168. PMID: 36977169; PMCID: PMC10055718.

20. Calasans TAS, Souza GTR, Melo CM, Madi RR, Jeraldo VLS. Socioenvironmental factors associated with *Schistosoma mansoni* infection and intermediate hosts in an urban area of northeastern Brazil. PLoS One. 2018;13(5):e0195519. doi: 10.1371/journal.pone.0195519.

21. Dabo A, Diarra AZ, Machault V, Touré O, Niambélé DS, Kanté A, Ongoiba A, Doumbo O. Urban Schistosomiasis and associated determinant factors among school children in Bamako, Mali West Africa. Infect Dis Poverty. 2015;4:4. 10.1186/2049-9957-4-4.

22. Diakité NR, Winkler MS, Coulibaly JT, Guindo-Coulibaly N, Utzinger J, N’Goran EK. Dynamics of freshwater snails and *Schistosoma* infection prevalence in schoolchildren during the construction and operation of a multipurpose dam in central Côte d’Ivoire. Infect Dis Poverty. 2017 May 4;6(1):93. doi: 10.1186/s40249-017-0305-3. PMID: 28468667; PMCID: PMC5415719.

23. Diarra AZ, Almeras L, Laroche M, Berenger JM, Koné AK, Bocoum Z, Dabo A, Doumbo O, Raoult D, Parola P. Molecular and MALDI-TOF identification of ticks and tick-associated bacteria in Mali. PLoS Negl Trop Dis. 2017;11(7):e0005762. doi: 10.1371/journal.pntd.0005762. PMID: 28742123; PMCID: PMC5542699.

24. Diaw OT, Vassiliades G. Epidemiology of schistosomiasis in livestock in Senegal. Rev Elev Med Vet Pays Trop. 1987;40(3):265–74.

25. Diaw OT, Vassiliades G, Seye M, Sarr Y. Prolifération de mollusques et incidence sur les trématodoses dans la région du delta et du lac de Guiers après la construction du barrage de Diama sur le fleuve Sénégal. Rev Elev Med Vet Pays Trop. 1990;43(4):499–502.

26. Ekin İ, Şeşen R, Alkan H, Akbal E, Başhan M. Protein content of five tissues from some edible and non-edible snails and bivalves distributed within Turkish territories. Int J Adv Biol Res. 2016;6(4):435–8.

27. Etard JF, Borel E. Contacts homme-eau et schistosomiase urinaire dans un village mauritanien [Man-water contacts and urinary schistosomiasis in a Mauritanian village]. Rev Epidemiol Sante Publique. 1992;40(4):268–275.

28. Folmer O, Black M, Hoeh W, Lutz R, Vrijenhoek R. DNA primers for amplification of mitochondrial cytochrome c oxidase subunit I from diverse metazoan invertebrates. Mol Mar Biol Biotechnol. 1994;3(5):294–9.

29. Gaye PM, Doucouré S, Sow D, Sokhna C, Ranque S. Identification of *Bulinus forskalii* as a potential intermediate host of *Schistosoma hæmatobium* in Senegal. PLoS Negl Trop Dis. 2023;17(5):e0010584.

30. Gaye PM, Ndiaye EHI, Doucouré S, Sow D, Gaye M, Goumballa N, Cassagne C, L’Ollivier C, Medianikov O, Sokhna C, Ranque S. Matrix-assisted laser desorption/ionization time-of-flight mass spectrometry traces the geographical source of Biomphalaria pfeifferi and *Bulinus forskalii*, involved in schistosomiasis transmission. Infect Dis Poverty. 2024;13(1):11. doi: 10.1186/s40249-023-01168-y.

31. Gaye PM, Doucoure S, Senghor B, Faye B, Goumballa N, Sembène M, L’Ollivier C, Parola P, Ranque S, Sow D, et al. *Bulinus Senegalensis* and *Bulinus Umbilicatus* Snail Infestations by the *Schistosoma haematobium* group in Niakhar Senegal. Pathogens. 2021;10:860. 10.3390/pathogens10070860

32. Gbalégba NGC, Silué KD, Ba O, Ba H, Tian-Bi NTY, Yapi GY, et al. Prevalence and seasonal transmission of *Schistosoma hæmatobium* infection among school-aged children in Kaedi town, southern Mauritania. Parasit Vectors. 2017;10(1):353.

33. Ghazy RM, Ellakany WI, Badr MM, Taktak NEM, Elhadad H, Abdo SM, Hagag A, Hussein AR, Tahoun MM. Determinants of *Schistosoma mansoni* transmission in hotspots at the late stage of elimination in Egypt. Infect Dis Poverty. 2022;11(1):102. doi: 10.1186/s40249-022-01026-3.

34. Greer GJ, Mimpfoundi R, Malek EA, Joky A, Ngonseu E, Ratard RC. Human schistosomiasis in Cameroon. II. Distribution of the snail hosts. Am J Trop Med Hyg. 1990;42(6):573–80. doi: 10.4269/ajtmh.1990.42.573. PMID: 2372088.

35. Gryseels B, Polman K, Clerinx J, Kestens L. Human schistosomiasis. Lancet. 2006;368(9541):1106–18. doi: 10.1016/S0140-6736(06)69440-3.

36. Hamlili FZ, Thiam F, Laroche M, Diarra AZ, Doucouré S, Gaye PM, et al. MALDI-TOF mass spectrometry for the identification of freshwater snails from Senegal, including intermediate hosts of schistosomes. PLoS Negl Trop Dis. 2021;15(9):e0009725.

37. Huguenin A, Depaquit J, Villena I, Ferté H. MALDI-TOF mass spectrometry: A new tool for rapid identification of cercariae (Trematoda, Digenea). Parasite. 2019.26. 11. 10.1051/parasite/2019011.

38. Jobin WR, Negrón-Aponte H, Michelson EH. Schistosomiasis in the Gorgol Valley of Mauritania. Am J Trop Med Hyg. 1976;25(4):587–94.

39. Joof E, Sanneh B, Sambou SM, Wade CM. Species diversity and distribution of schistosome intermediate snail hosts in The Gambia. PLoS Negl Trop Dis. 2021b; 15(10): e0009823. 10.1371/journal.pntd.0009823

40. Kane R, Stothard JR, Emery AM, Rollinson D. Molecular characterization of freshwater snails in the genus *Bulinus*: A role for barcodes?. Parasites & vectors. 2008 ;1(1)15. 10.1186/1756-3305-1-15.

41. Kim TS, Pak JH, Kim JB, Bahk YY. Clonorchis sinensis, an oriental liver fluke, as a human biological agent of cholangiocarcinoma: a brief review. BMB Rep. 2016 Nov;49(11):590–597. doi: 10.5483/bmbrep.2016.49.11.109.

42. Kumar S, Stecher G, Tamura K. MEGA7: Molecular Evolutionary Genetics Analysis Version 7.0 for Bigger Datasets. Mol Biol Evol. 2016;33(7):1870–1874. doi: 10.1093/molbev/msw054.

43. Labbo R, Djibrilla A, Zamanka H, Garba A, Chippaux JP. *Bulinus forskalii*: a new potential intermediate host for *Schistosoma hæmatobium* in Niger. Trans R Soc Trop Med Hyg. 2007;101(8):847–8. doi: 10.1016/j.trstmh.2007.03.016.

44. Lafri I, Almeras L, Bitam I, Caputo A, Yssouf A, Forestier C-L, et al. Identification of Algerian Field Caught Phlebotomine Sand Fly Vectors by MALDI-TOFMS.PLoSNeglTropDis.2016;10:e0004351. 10.1371/journal.pntd.0004351 PMID: 26771833

45. Léger E, Borlase A, Fall CB, Diouf ND, Diop SD, Yasenev L, Catalano S, Thiam CT, Ndiaye A, Emery A, et al. Prevalence and distribution of Schisto somiasis in human, livestock, and snail populations in Northern Senegal: a one health epidemiological study of a multi-host system. Lancet Planetary Health. 2020;4:E330–42.

46. Marill FG. Diffusion de la bilharziose chez les bovins, ovins et caprins en Mauritanie et dans la vallée du Sénégal. Bull Acad Natl Méd. 1961a;145(6):147–50.

47. Marill FG. Enseignements d’une première enquête sur l’épidémiologie de la bilharziose à *Schistosoma hæmatobium* en Mauritanie [The epidemiology of urinary schistosomiasis in Mauritania. A preliminary study.]. Med Trop (Mars). 1961b;21:373–86.

48. Nakatt L, Gaye PM, Moukah MO, Niang B, Basco L, Ranque S, Ould Mohamed Salem Boukhary A. Urogenital schistosomiasis in schoolchildren in the lake zones of Kankossa and Oued Rawdha, southern Mauritania: The first parasitological and malacological survey. PLoS Negl Trop Dis. 2024;18(9):e0012505. doi: 10.1371/journal.pntd.0012505. PMID: 39321164; PMCID: PMC11458011.

49. Nebbak A, El Hamzaoui B, Berenger JM, Bitam I, Raoult D, Almeras L, Parola P. Comparative analysis of storage conditions and homogenization methods for tick and flea species for identification by MALDI-TOF MS. Med Vet Entomol. 2017;31(4):438–448. doi: 10.1111/mve.12250.

50. Nebbak A, Monteil-Bouchard S, Berenger J-M, Almeras L, Parola P, Desnues C. Virome diversity among mosquito populations in a sub-urban Region of Marseille, France. Viruses. 2021;13:768.

51. Ngoy S, Diarra AZ, Laudisoit A. et al. Using MALDI-TOF mass spectrometry to identify ticks collected on domestic and wild animals from the Democratic Republic of the Congo. Exp Appl Acarol. 2021;84:637–657. 10.1007/s10493-021-00629-z

52. Oso OG, Odaibo AB. Land use/land cover change, physico-chemical parameters and freshwater snails in Yewa North, Southwestern Nigeria. PLoS One. 2021;16(2):e0246566. 10.1371/journal.pone.0246566 PMID: 33556093

53. Ouldabdallahi M, Ouldbezeid M, Diop C, Dem E, Lassana K. Epidemiology of human schistosomiasis in Mauritania. The right bank of the Senegal River as model. Bull Soc Pathol Exot. 2010;103(5):317–22.

54. Ould Ahmed Salem CB, Boussery A, Hafid J. Study of prevalence and parasite load of *Schistosoma hæmatobium* in schoolchildren in the Rosso region, Mauritania. Med Sante Trop. 2019;29(3):268–72.

55. Pinto HA., De Melo AL. A checklist of trematodes (Platyhelminthes) transmitted by *Melanoides tuberculata* (Mollusca: Thiaridae). Zootaxa.2011;2799(2799):15–28. 10.11646/zootaxa.2799.1.2.

56. Raahauge P, Kristensen TK. A comparison of *Bulinus* africanus group species (Planorbidae; Gastropoda) by use of the internal transcribed spacer 1 region combined by morphological and anatomical characters. Acta Trop. 2000;75(1):85–94. doi: 10.1016/s0001-706x(99)00086-8. PMID: 10708010.

57. Sène AM. Développement durable et impacts des politiques publiques de gestion de la vallée du fleuve Sénégal : du régional au local. [VertigO] La revue électronique en sciences de l’environnement. 2009;9(3):17. doi : https://id.erudit.org/iderudit/044189ar.

58. Sène M, Southgate VR, Vercruysse J. *Bulinus truncatus*, hôte intermédiaire de *Schistosoma hæmatobium* dans le bassin du fleuve Sénégal [*Bulinus truncatus*, intermediate host of *Schistosoma hæmatobium* in the Senegal River Basin (SRB)]. Bull Soc Pathol Exot. 2004;97(1):29–32.

59. Senghor B, Diaw OT, Doucoure S, Seye M, Talla I, Diallo A, et al. Study of the snail intermediate hosts of urogenital schistosomiasis in Niakhar, region of Fatick, West central Senegal. Parasit Vectors. 2015;8:410.

60. Senghor B, Webster B, Pennance T, Sène M, Doucouré S, Sow D, et al. Molecular characterization of schistosome cercariae and their *Bulinus* snail hosts from Niakhar, a seasonal transmission focus in central Senegal. Curr Res Parasitol Vector Borne Dis. 2023;3:100114.

61. Sokouri EA, Ahouty B, Abé IA, Yao FGD, Konan TK, Nyangiri OA, MacLeod A, Matovu E, Noyes H, Koffi M; TrypanoGEN+ Research Group of the H3Africa Consortium. Evaluation of the epidemiological situation of intestinal schistosomiasis using the POC-CCA parasite antigen test and the Kato-Katz egg count test in school-age children in endemic villages in western Côte d’Ivoire. Parasite. 2024;31:66. doi: 10.1051/parasite/2024049.

62. Stephan R, Johler S, Oesterle N, Näumann G, Vogel G, Pflüger V. Rapid and reliable species identification of scallops by MALDI-TOF mass spectrometry. Food Control. 2014;46:6–9.

63. Strong EE, Gargominy O, Ponder WF, Bouchet P. Global diversity of gastropods (Gastropoda; Mollusca) in freshwater. Freshwater Animal Diversity Assessment. 2008;595:149–166. DOI 10.1007/S10750-007-9012-6.

64. Sunday JO & Oso OG. Environmental influences on the abundance and distribution of gastropods in Edu Local Government Area, North Central Nigeria. PLOS Water. 2025;4(9): e0000364. 10.1371/journal.pwat.0000364

65. Ugbomoiko US, Kareem II, Awe DO, Babamale AO, Gyang PV, Nwafor TE, Akinwale OP. Characterization of freshwater snail intermediate hosts of Schistosomes in four communities from Osun State, Southwest Nigeria. One Health Implement Res. 2022;2:88–95.

66. Tang ZL, Huang Y, Yu XB. Current status and perspectives of *Clonorchis sinensis* and clonorchiasis: epidemiology, pathogenesis, omics, prevention and control. Infect Dis Poverty. 2016;5(71). 10.1186/s40249-016-0166-1.

67. Tamura K, Nei M. Estimation of the number of nucleotide substitutions in the control region of mitochondrial DNA in humans and chimpanzees. Mol Biol Evol. 1993;10(3):512–526. doi: 10.1093/oxfordjournals.molbev.a040023.

68. Tian-Bi YNT, Webster B, Konan CK, et al. Molecular characterization and distribution of *Schistosoma* cercariae collected from naturally infected bulinid snails in northern and central Côte d’Ivoire. Parasites Vectors. 2019;12:117. 10.1186/s13071-019-3381-3.

69. Trippler L, Knopp S, Welsche S, Webster BL, Stothard JR, Blair L, Allan F, Ame SM, Juma S, Kabole F, Ali SM, Rollinson D, Pennance T. The long road to schistosomiasis elimination in Zanzibar: A systematic review covering 100 years of research, interventions and control milestones. Adv Parasitol. 2023;122:71–191. doi: 10.1016/bs.apar.2023.06.001.

70. Tumwebaze I, Clewing C, Dusabe MC, Tumusiime J, Kagoro-Rugunda G, Hammoud C, Albrecht C. Molecular identification of *Bulinus* Spp. Intermediate Host Snails of *Schistosoma* Spp. in Crater Lakes of Western Uganda with Implications for the transmission of the *Schistosoma hæmatobium* group parasites. Parasites Vectors. 2019;12(565). 10.1186/s13071-019-3811-2.

71. Wang Y, Zhang X, Wang X, Zhang N, Yu Y, Gong P, Zhang X, Ma Y, Li X, Li J. *Clonorchis sinensis* aggravates biliary fibrosis through promoting IL-6 production via toll-like receptor 2-mediated AKT and p38 signal pathways. PLoS Negl Trop Dis. 2023 Jan 24;17(1):e0011062. doi: 10.1371/journal.pntd.0011062. PMID: 36693049; PMCID: PMC9873171.

72. Wepnje GB, Peters MK, Green AE, Nkuizin TE, Kenko DBN, Dzekashu FF, et al. Seasonal and environmental dynamics of intra-urban freshwater habitats and their influence on the abundance of *Bulinus* snail host of *Schistosoma hæmatobium* in the Tiko endemic focus, Mount Cameroon region. PLoS One. 2023;18(10):e0292943. 10.1371/journal.pone.0292943 PMID: 37856526

73. WHO. WHO Guideline on Control and Elimination of Human Schistosomiasis. World Health Organization: Geneva, Switzerland. 2022. Accessible en ligne: https://www.who.int/publications/i/item/9789240041608. Consulté le 15 Mars 2024.

74. Wright CA. Taxonomic problems in the molluscan genus *Bulinus*. Trans R Soc Trop Med Hyg. 1961;55:225–31.

75. Yssouf A, Parola P, Lindström A, Lilja T, L’Ambert G, Bondesson U, Berenger JM, Raoult D, Almeras L. Identification of European mosquito species by MALDI-TOF MS. Parasitol Res. 2014;113(6):2375–2378. doi: 10.1007/s00436-014-3876-y.

76. Zein-Eddine R, Djuikwo-Teukeng FF, Al-Jawhari M, Senghor B, Huyse T, Dreyfuss G. Phylogeny of seven *Bulinus* species originating from endemic areas in three African countries, in relation to the human blood fluke *Schistosoma hæmatobium*. BMC Evol Biol. 2014;14:271. doi: 10.1186/s12862-014-0271-3.

77. Zurita A, Djeghar R, Callejón R, Cutillas C, Parola P, Laroche M. Matrix-assisted laser desorption/ionization time-of-flight mass spectrometry as a useful tool for the rapid identification of wild flea vectors preserved in alcohol. Med Vet Entomol. 2019;33(2):185–194. doi: 10.1111/mve.12351. Epub 2018 Dec 5. PMID: 30516832.

78. Zongo D, Kabre BG, Dayeri D, Savadogo B, Poda J-N. Parasitological profile of two forms of schistosomiasis (urinary and intestinal forms) at ten sites in Burkina Faso (Sub-Saharan Africa country). C R Biol. 2013;336:317–319. 10.1016/j.crvi.2013.04.014

